# The role of matrilineality in shaping patterns of Y chromosome and mtDNA sequence variation in southwestern Angola

**DOI:** 10.1101/349878

**Authors:** Sandra Oliveira, Alexander Hübner, Anne-Maria Fehn, Teresa Aço, Fernanda Lages, Brigitte Pakendorf, Mark Stoneking, Jorge Rocha

**Author notes:** **Corresponding author** Sandra Oliveira, CIBIO/InBIO, Campus de Vairão, Rua Padre Armando Quintas n 7, 4485-661 Vairão, Portugal.

## Abstract

Southwestern Angola is a region characterized by contact between indigenous foragers and incoming food-producers, involving genetic and cultural exchanges between peoples speaking Kx’a, Khoe-Kwadi and Bantu languages. Although present-day Bantu-speakers share a patrilocal residence pattern and matrilineal principle of clan and group membership, a highly stratified social setting divides dominant pastoralists from marginalized groups that subsist on alternative strategies and have previously been though to have pre-Bantu origins. Here, we compare new high-resolution sequence data from 2.3 Mb of the non-recombining Y chromosome (NRY) from 170 individuals with previously reported mitochondrial genomes (mtDNA), to investigate the population history of seven representative southwestern Angolan groups (Himba, Kuvale, Kwisi, Kwepe, Twa, Tjimba, !Xun) and to study the causes and consequences of sex-biased processes in their genetic variation. We found no clear link between the formerly Kwadi-speaking Kwepe and pre-Bantu eastern African migrants, and no pre-Bantu NRY lineages among Bantu-speaking groups, except for small amounts of “Khoisan” introgression. We therefore propose that irrespective of their subsistence strategies, all Bantu-speaking groups of the area share a male Bantu origin. Additionally, we show that in Bantu-speaking groups, the levels of among-group and between-group variation are higher for mtDNA than for NRY. These results, together with our previous demonstration that the matriclanic systems of southwestern Angolan Bantu groups are genealogically consistent, suggest that matrilineality strongly enhances both female population sizes and interpopulation mtDNA variation.

## Introduction

Due to their uniparental modes of inheritance, the mtDNA and NRY have been extensively used to study differences between paternal and maternal histories of human populations and assess the influence of socio-cultural practices on sex-specific patterns of variation ^1,2^. However, direct comparisons of mtDNA and NRY diversity were hampered until recently by differences between methods used to detect variation, and by ascertainment bias in the choice of NRY single nucleotide polymorphisms (SNPs) ^3^. In the last few years, various studies took advantage of the increasing availability of next generation sequencing (NGS) platforms to compare unbiased NRY and mtDNA sequence data on a global scale ^4–7^. Nevertheless, the sex-specific patterns disclosed by worldwide studies reflect average trends exhibited by different macro-regions, and do not explore the rich diversity of interactions between genetic variation and cultural practices that shape NRY and mtDNA variation at the local level ^8^.

Here, we used targeted NGS to generate unbiased sequence data from 2.3 Mb of NRY in 170 males comprising seven small communities from SW Angola, whose complete mtDNA genomes have been previously investigated ^9^. Despite being located in a relatively small geographic area (Fig. 1), these groups offer a unique framework to explore the relationships between socio-cultural practices, local variation in mtDNA versus NRY diversity, and continent-wide migratory processes. The !Xun from Kunene Province are Kx’a-speaking hunter gatherers who descend from the oldest population layer of southern Africa represented by the so-called “Khoisan” peoples ^10^. The Kuvale, Himba, Tjimba, Twa, Kwisi and Kwepe from the Angolan Namib Desert are all Bantu-speaking groups, who display striking socio-economic disparities that have been associated with different population histories. The Himba and Kuvale are two mildly polygynous pastoralist populations belonging to the broad Herero ethnic division, who arrived in SW Africa during the Bantu expansions ^11^. The Kwepe, Twa, Kwisi and Tjimba are marginalized ethnic minorities subsiding on small-scale pastoralism and foraging, who gravitate around the Himba and Kuvale and are best described as peripatetic peoples ^12^. While the Himba-speaking Tjimba are commonly thought to be impoverished Himba ^13^, the Kuvale-speaking Kwisi and Twa have been associated with a hypothetical stratum of pre-Bantu foragers of unknown provenance, whose original language has been lost ^14^. The Kwepe, who spoke Kwadi – a now extinct language belonging to the Khoe-Kwadi family – before shifting to Kuvale, were considered to be remnants of a pre-Bantu pastoralist migration introducing Khoe-Kwadi languages into southern Africa ^15^. In spite of their different subsistence strategies, all Bantu-speaking groups of the Angolan Namib share a patrilocal residence pattern and a matrilineal descent-group system, which regulates important parts of social life, such as group membership, inheritance and marriage behaviour.

**Figure 1.**
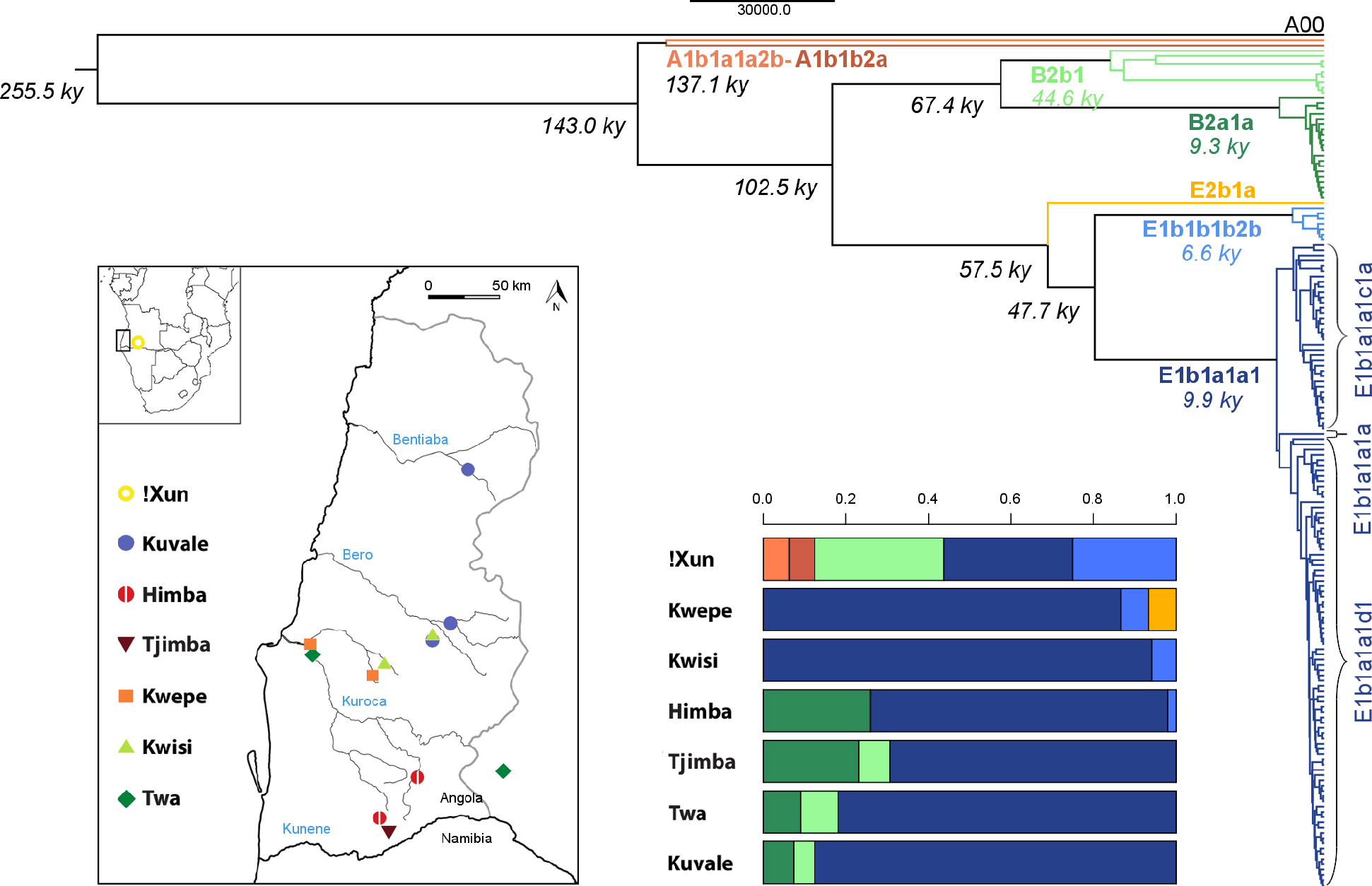
Y chromosome phylogeny, haplogroup distribution and map of the sampling locations. The phylogenetic tree was reconstructed in BEAST based on 2,379 SNPs and is in accordance with the known Y chromosome topology. Main haplogroup clades and their labels are shown with different colors. Age estimates are reported in italics near each node, with the TMRCA of main haplogroups shown with their corresponding color. A map of the sampling locations, re-used with permission from Oliveira et al. (2018) ^9^, is shown on the bottom left, and the haplogroup distribution per population is shown on the bottom right, with color-coding corresponding to the phylogenetic tree.

In this study, we compare new data on NRY sequence variation with our previous findings on whole mtDNA genomes to investigate whether the histories of these groups fit the expectations of earlier anthropological and linguistic hypotheses, and to study the causes and consequences of sex-biased processes for their genetic variation.

## Material and Methods

### Samples

We analyzed 162 partial NRY sequences sampled from populations that inhabit the Angolan Namib Desert (including the Himba, Kuvale, Kwepe, Kwisi, Twa and Tjimba) and the Kunene Province (!Xun) (Fig. 1; Table S1), and eight additional sequences from Bantu-speaking individuals with other ethnic affiliations used exclusively in haplotype-based analysis. The samples were collected as described previously ^16^, with the donors’ written informed consent, the ethical clearance of ISCED and CIBIO/InBIO-University of Porto boards, and the support and permission of the Provincial Governments of Namibe and Kunene.

### NRY Sequencing

Indexed libraries produced previously ^9^ were enriched for 2.3 Mb of target NRY as previously described ^17^. Paired-end sequencing data of 100+7 bp length were generated on the Illumina HiSeq 2500 platform and standard Illumina base-calling was performed using Bustard. We trimmed Illumina adapters and merged completely overlapping paired sequences using leeHOM ^18^, and de-multiplexed the pooled sequencing data using deML ^19^. The sequencing data were aligned to the human reference genome hg19 and SNPs were identified following ^17^. Y chromosome haplogroups were determined using yhaplo ^20^, with two modifications: i) the SNP defining B2b1 was corrected according to a recent ISOGG update (August 6th, 2017); and ii) since the mutation commonly used to identify haplogroup E1b1a1a1c1a1 was not typed in this study (P252/U174), we based our E1b1a1a1c1a1 assignment on the presence of mutation V1245/M4671, previously identified on E1b1a1a1c1a1 branches ^21,22^.

### Data analysis

Genetic diversity indices, pairwise Фst values and Analyses of Molecular Variance (AMOVA) were computed in Arlequin v35 (ref. 23). Non-metric multidimensional scaling (MDS), k-means and neighbor-joining (NJ) analyses based on pairwise Фst distance matrices were carried out in R, using the functions “isoMDS”, “kmeans” with several random starts, and “nj”, respectively. To determine the support of NJ partitions we generated bootstrap replicates with the function “boot.phylo” and used “stat.phist” (strataG v0.9.2) to recalculate Фst distances.

A phylogenetic tree was constructed with BEAST v1.8 (ref. 24), using an A00 representative haplotype as outgroup ^4^. To account for the absence of invariable sites in BEAST, we applied an invariant site correction. We used a strict clock and a mutation rate of 0.74×10^−9^ mutations/bp/year, as estimated by Karmin et al. (2015) ^4^ based on calibration with two ancient DNA sequences. Additional settings used in BEAST are reported in Table S2. We performed additional BEAST runs to build Bayesian Skyline plots (BSP) for different population groupings based on NRY and previously published mtDNA data ^9^, and also for specific NRY haplogroups (see settings in Table S2).

Median-joining networks were computed with Network 5.0 (www.fluxus-engineering.com) and plotted with Network Publisher v2.1.1.2.

For comparative purposes, we merged the sequence data generated in this study (2.3Mb) with i) 447 partial NRY sequences (0.9Mb) from other southern African groups ^25^, obtaining an overlap of 0.56 Mb, and ii) 21 complete Y chromosomes of various origins in Africa ^22,26^. The merged datasets were used to build networks.

## Results

We obtained 2.3 Mb of NRY sequence from 170 Angolan individuals, with a mean coverage of 28x (range 8-52x). After quality filtering, a total of 1854 SNPs were identified, of which only 66% are reported in dbSNP (build 150). A VCF file containing all SNPs and 154 non-variable nucleotide positions that are different from the reference sequence is available online (Supplementary datafile 1). An average of 6 nucleotides per individual (0.3%) were missing and were imputed with Beagle 4.0 (ref. 27).

### NRY phylogeography in Angola

Figure 1 displays a Bayesian phylogenetic tree for the Angolan NRY sequences, which also includes an early splitting A00 haplotype ^4^(see also Fig. S1 for a network relating all Angolan haplotypes). The estimated split time of the A1b1 branch (143 kya), which corresponds to the most recent common ancestor (TMRCA) of all Angolan sequences, and their split from A00 (256 kya), are remarkably close to previous estimates based on high-coverage whole Y chromosomes sampled from other populations (Table S3) ^4,28^. Despite being sampled in a relatively small area, the Angolan lineages have very different phylogeographical characteristics, and belong to haplogroups that have been associated with three major population layers settling southern Africa at different periods (see Table S4 for alternative haplogroup nomenclatures). A1b1a, A1b1b and B2b contain deep-rooting nodes and are associated with an early substrate of “Khoisan” foragers (>10 kya) speaking Kx’a and Tuu languages ^25^. These represent 44% of the ! Xun but only 0-9% of the genetic makeup of Bantu-speaking peoples from Angola (Fig. 1; Table S4). E1b1b sequences, which have been linked to a pre-Bantu migration of sheep pastoralists from East Africa (~2 kya) ^29^ have a TMRCA of 6.6 kya and are observed in varying frequencies among the Kwepe (7%), Kwisi (6%), Himba (2%) and !Xun (25%) (Fig. 1; Table S4). E1b1a and B2a, which have been previously associated with the Bantu expansions ^30,31^(though B2a might also have existed in Khoisan groups before the arrival of Bantu speakers ^25^), have TMRCAs close to 10 kya and represent 91-98% of the NRY sequences sampled among Bantu-speaking groups and 33% of the !Xun NRY (Fig. 1; Table S4). In accordance with previous studies ^7^, subhaplogroup E1b1a1a1 displays a star-like branching pattern (Figs. 1 and S1), consistent with a rapid demographic expansion from a small ancestral population size. This is also observed in the population-based BSPs from Bantu speaking groups (Fig. 2) and in the lineage-specific BSPs, which display strong population growth in E1b1a1a1 and its subclades (E1b1a1a1d1, E1b1a1a1c1a), as well as an additional sign of moderate growth in B2a1a (Fig. S2).

**Figure 2.**
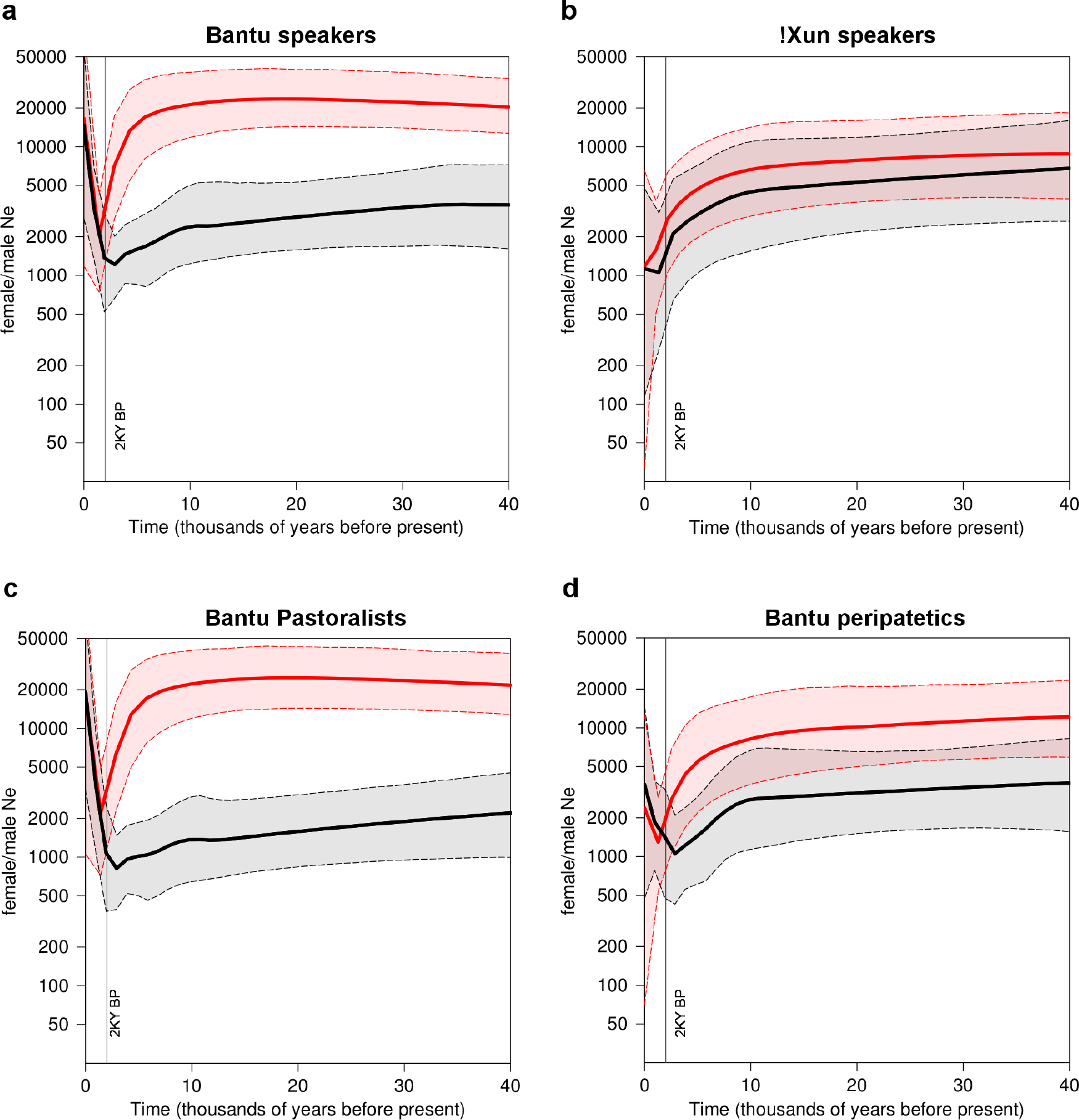
Bayesian skyline plots (BSP) of effective population size change through time, based on mtDNA (red) and the NRY (black). Thick lines show the mean estimates and dashed lines show the 95% HPD intervals. The vertical line highlights the 2 ky before present mark. Effective sizes are plotted on a log scale. Generation times of 25 and 31 years were assumed for mtDNA and the NRY, respectively ^32^.

An inspection of the molecular relationships between NRY haplotypes from different populations reveals that most lineages from Angola cluster together with other available sequences from southern Africa (Fig. S3). The only exceptions are the B2a1a sequences (Fig. S3g-h), which are grouped in a divergent monophyletic cluster that includes lineages previously found in SW Bantu groups from Namibia, and the B2b1 haplotypes, which are very divergent from haplotypes found in the wider region of southern Africa or in Pygmy groups (Fig. 3c-d).

**Figure 3.**
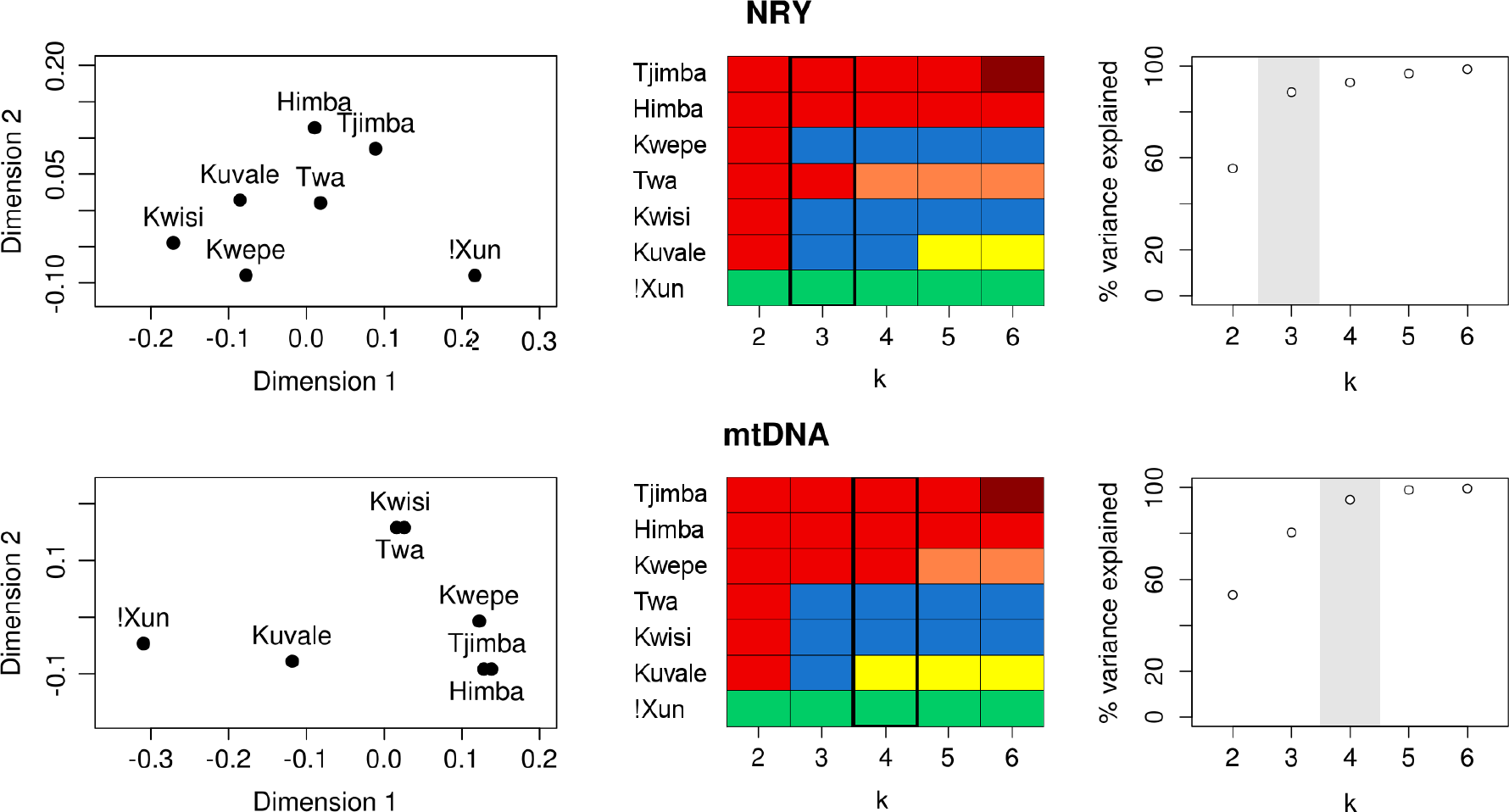
Multidimensional scaling (MDS) and k-means analysis based on Фst distances of mtDNA and the NRY. a) MDS. The stress values are 0.003 and 0.006 for NRY and mtDNA, respectively. b) k-means. Each color represents the cluster assigned to a population. c) Percentage of variance explained by each k. The best k (i.e. the point after which the variance explained has diminishing returns) is highlighted in grey.

### Intrapopulation diversity and demographic inferences

Table S1 presents summary statistics for NRY and mtDNA diversity. The NRY nucleotide diversity in the !Xun (π_NRY_ = 1.2 × 10^−4^) is 2.5 times higher than in the Bantu-speaking groups (π_NRY_ = 4.8 × 10^−5^), and similar to values calculated from previous studies for other “Khoisan” groups (π_NRY_ = 1.5 × 10^−4^ − 1.8 × 10^−4^) ^6,25^.

In Bantu speakers, NRY vs. mtDNA diversity ratios accounting for differences in mutation rates of the two chromosomes (π_NRY_/ π_mtDNA_) range from 0.09 to 0.5 (Table S1), indicating that Bantu peoples, like many other human populations, display less NRY diversity relative to mtDNA than expected in neutral demographic models without sex-biased processes ^4,6,25,33^. In contrast, the !Xun resemble other “Khoisan” groups in displaying comparable levels of diversity in both sexes (π_NRY_/ π_mtDNA_ = 1.11) (see ^6,25^).

To better understand the present differences in levels of mtDNA and NRY diversity, we inferred the history of male and female effective population size (*Ne*) changes by using BSPs (Fig. 2). We found striking population size differences between males and females in a pooled sample comprising all Bantu-speaking groups from the Angolan Namib (Fig. 2a). Starting from the past, *Ne* estimates based on mtDNA (*Ne*_f_) remained stable for a long period of time (~20,000), and display a sharp reduction with minimum size (~2,000) around 2 kya, followed by expansion to the present (Fig. 2a). In contrast, the male demographic profile is characterized by a recent expansion from a relatively low, more stable long-term population size (*Ne*_m_ ~3,000) (Fig. 2a). Unlike the Bantu-speaking populations, the !Xun displayed almost overlapping female and male population sizes that start to decline around 10 kya with no traces of population recovery (Fig 2b).

When population size changes among Bantu speakers with different subsistence patterns were compared, the peripatetic communities showed less pronounced differences in sex-specific *Ne*, and smaller post-bottleneck size recoveries than the pastoral populations (Fig. 2c-d). These differences persisted when the lower sample sizes of peripatetics were taken into account (Fig. S4).

As BSPs assume a single, isolated, panmictic population, and the Angolan groups are likely to be part of a network of structured populations, some inferred demographic events might have been more influenced by migration levels and sampling design than by real changes in population size ^34^. To account for these confounding factors, we generated separate mtDNA and NRY BSPs for all individual groups (Fig. S5). The demographic profile of the Kuvale, who display high frequencies of “Khoisan”-related mtDNA haplogroups, remained similar to the Himba, who have similar sample sizes and do not show signs of “Khoisan” introgression in their mtDNA ^9^, suggesting that the differences between female and male BSPs of pastoralists are not exclusively due to admixture with resident foragers (Fig. S5). On the other hand, not all of the BSPs for individual peripatetic groups show the signs of post-bottleneck *Ne* recovery that were detected in the pooled “peripatetic” sample (Fig. S5).

### Interpopulation diversity

To compare the levels of between-population divergence for the NRY with our previous results on mtDNA variation in the same populations ^9^, we carried out AMOVA based on different partitions of the data (Table S5). Although we found similar amounts of divergence between the !Xun and Bantu speakers (22.5% NRY vs. 16.6% mtDNA), the genetic differentiation among Bantu speakers is much lower for the NRY than for mtDNA (4.4% NRY vs. 20.2% mtDNA), even when the Kuvale are removed from the comparisons to eliminate the confounding effects of “Khoisan” lineages on the levels of population divergence (5.5% NRY vs. 18.8% mtDNA) ^9^(Table S5). Moreover, we found that the genetic differentiation among matriclans for mtDNA (50.8%) is much higher than for the NRY (2.5%) (Table S5), reflecting the structuring effect of the matriclanic system on mtDNA, but not on NRY variation (see ^9^).

The population relationships displayed in an MDS plot based on pairwise Фst values further reveal noticeable differences between the NRY and mtDNA (Fig. 3a), which are reflected in a lack of correlation between their corresponding Фst matrices (Mantel test, p-value = 0.091). In addition, there is a clear mismatch between the clustering patterns inferred from k-means analyses based on NRY and mtDNA (Fig. 3b-c). For mtDNA, the best k-means partition (k = 4; Фct = 20.9%; p-value < 0.006; Table S5) places the Kwisi and Twa in a separate group, and associates the Kwepe with the Himba. For NRY (k = 3; Фst = 14.4%; p-value < 0.014; Table S5), the Kwepe and the Kwisi are grouped with their northern Kuvale neighbors, while the southernmost Twa are grouped with the Himba (Figs. 1 and 3). Interestingly, the NRY clustering has remarkable parallels with the distribution of cultural traits like language, dressing habits, and names of matriclans. For example, while the Twa tend to imitate the dressing habits of Himba women, the Kwisi and Kwepe try to mimic the characteristic attire of the Kuvale ^14^. Moreover, the variety of Kuvale spoken by the Twa has clearly been influenced by the Himba language, while the Kwisi and Kwepe speak language varieties that are practically indistinguishable from mainstream Kuvale ^9^(see NJ in Fig. S6a). This is also reflected in the significant correlation we found between lexicon-based linguistic distances ^9^ and NRY distances (Mantel test, p-value = 0.033). Finally, we have previously shown that peripatetic groups tend to replace their own clan names with those of their neighboring pastoral groups, leading to the shared use of matriclan labels by the Twa and Himba on the one hand, and by the Kwisi, Kwepe and Kuvale on the other ^9^. Despite the genetic consistency of the matriclanic system within each group, this clan switching leads to quite different patterns of population relationships based on mtDNA variation and on the distribution of clan names (Fig. 6b-c). In contrast, we found that NJ trees constructed based on Фst distances for NRY and on distances based on clan name frequencies do have similar patterns (Fig. S6b, d), as confirmed by a significant correlation between the two distance matrices (Mantel test, p-value = 0.001).

## Discussion

### The origins of NRY diversity in SW Angola

In accordance with our previous mtDNA study ^9^, the present NRY analysis reveals a major division between the Kx’a-speaking !Xun and the Bantu-speaking groups, whose paternal genetic ancestry does not display any old remnant lineages, or a clear link to pre-Bantu eastern African migrants introducing Khoe-Kwadi languages and pastoralism into southern Africa (cf. ^15^). This is especially evident in the distribution of the eastern African subhaplogroup E1b1b1b2b ^29^, which reaches the highest frequency in the !Xun (25%) and not in the formerly Kwadi-speaking Kwepe (7%). This observation, together with recent genome-wide estimates of 9-22% of eastern African ancestry in other Kx’a and Tuu-speaking groups ^35^, suggests that eastern African admixture was not restricted to present-day Khoe-Kwadi speakers. Alternatively, it is likely that the dispersal of pastoralism and Khoe-Kwadi languages involved a series of punctuated contacts that led to a wide variety of cultural, genetic and linguistic outcomes, including possible shifts to Khoe-Kwadi by originally Bantu-speaking peoples ^36^.

Although traces of an ancestral pre-Bantu population may yet be found in autosomal genome-wide studies, the extant variation in both uniparental markers strongly supports a scenario in which all groups of the Angolan Namib share most of their genetic ancestry with other Bantu groups but became increasingly differentiated within the highly stratified social context of SW African pastoral societies ^11^.

### The influence of socio-cultural behaviors on the diversity of NRY and mtDNA

A comparison of the NRY variation with previous mtDNA results for the same groups ^9^identifies three main sex-specific patterns. First, gene flow from the Bantu into the !Xun is much higher for male than for female lineages (31% NRY vs. 3% mtDNA), similar to the reported male-biased patterns of gene flow from Bantu to Khoisan-speaking groups ^33^, and from non-Pygmies to Pygmies in Central Africa ^37^. A comparable trend, involving exclusive introgression of NRY eastern African lineages into the !Xun (25%) was also found. These patterns may be explained by a context of social discrimination, in which women from food producing populations are prevented from moving into forager communities, while food producing men and forager women can have children, who will then mostly be raised in the mother’s group ^37,38^. However, the dominant Kuvale pastoralists, who show a high frequency of “Khoisan”-related mtDNA (53%), indicate that admixed children may also remain in the father’s group ^9^.

Secondly, the levels of intrapopulation diversity in the Bantu-speaking peoples from the Namib were found to be consistently higher for mtDNA than for the NRY, reflecting the marked association between the Bantu expansion and the relatively young NRY E1b1a1a1 haplogroup, which has no parallel in mtDNA ^25,39^. In contrast, the !Xun have a more diverse NRY haplogroup composition, combining deeply rooted lineages and younger clades obtained through recent admixture. Using BSP analysis, we found these patterns to be reflected in larger long-term *Ne*_f_ than *Ne*_m_ in Bantu speakers, and more equal sex-specific *Ne* in the !Xun (Figs. 2 and S3).

Global patterns showing that *Ne*_f_ was larger than *Ne*_m_ during a large part of human history have been explained by a number of sex-biased processes, including natural selection affecting the NRY, or culturally influenced sex-specific demographic behaviors ^1–3^. In the context of the Bantu expansions, these patterns have been mostly interpreted as the result of polygyny and/or higher levels of assimilation of females from resident forager communities ^38,40^. However, most groups from the Angolan Namib are only mildly polygynous ^11^and ethnographic data suggest that the actual rates of polygyny in many populations may be insufficient to significantly reduce *Ne*_*m*_ ^2,41^. In addition, the finding of a large *Ne*_f_/*Ne*_m_ ratio in the Himba (Fig. S5), who have almost no Khoisan-related mtDNA lineages ^9^, indicates that female biased introgression cannot fully explain the observed patterns.

An alternative explanation may be sought in the prevailing matrilineal descent rules, which might have created a sex-specific structuring effect, similar to that proposed for patrilineal groups from Central Asia ^42^. As we previously demonstrated ^9^, all Bantu-speaking groups sampled in the region have genetically consistent, highly structured matrilineal descent systems, with levels of genetic variation between matriclans that are 20 times higher for mtDNA than for the NRY (50.8% mtDNA vs. 2.5% NRY; Table S5). Since ethnic groups are conglomerates of matriclans, they harbor a remarkable amount of mtDNA structure and have fragmented female populations that can inflate *Ne*_*f*_ estimates ^43^. Under this hypothesis, and using the terminology proposed by Wakeley (1999), the population size growth starting at ~2 kya that is detected in both the female and male BSPs (Figs. 2 and S5) would be associated with the “scattering phase” of the mtDNA tree. The separation of male and female BSPs before 2 kya would then correspond to Wakeley’s “collecting phase” and reflect the inflation of *Ne*_*f*_ due to the large mtDNA differences between matriclans. Since the male pool is not structured, the NRY tree has no collecting phase and *Ne*_*m*_ remains essentially unchanged beyond 2 kya. In the future, it will be interesting to make a more comprehensive re-evaluation of the relationship between descent rules and *Ne*_*m*_/*Ne*_*f*_ ratios across different Bantu populations, since studies in other regions of the world have shown that the more structured sex may not display the highest *Ne* if the extinction rate of clans is high ^42,44^. The third important sex-specific pattern observed in this study is the much lower amount of between-group differentiation for NRY than for mtDNA among Bantu-speaking populations (4.4% NRY vs. 20.2% mtDNA), in spite of the patrilocal residence patterns of all ethnic groups (Table S5). This difference can hardly be explained by unequal levels of introgression of “Khoisan” mtDNA lineages into the Bantu, since the percentage of mtDNA variation remains high (18.8%) when the Kuvale, who have high frequencies of “Khoisan”-related mtDNA, are excluded from the comparisons. It therefore seems more plausible that differentiation is higher in the mtDNA simply because there is more ancestral mtDNA than NRY variation that can be sorted among different populations (see ^45^). Moreover, due to the matriclanic organization of all Bantu-speaking communities, factors enhancing inter-group differentiation, like kin-structured migration and kin-structured founder effects ^46^, would have been restricted to mtDNA. Finally, it is also likely that the discrepancy between among-group divergence of mtDNA and NRY might have been influenced by higher migration rates in males than females. In fact, although all Bantu-speaking populations have patrilocal residence patterns, the observance of endogamy rules severely constrains the between-group mobility of females. In this context, the children from extramarital unions involving members from different populations tend to be raised in the mother’s group, effectively increasing male versus female migration rates. Moreover, it is likely that, in the highly hierarchized setting of the Namib, most intergroup extramarital unions would involve men from dominant groups and women from peripatetic communities. This hypothesis is indirectly supported by the finding that in NRY-based clusters (but not in mtDNA) pastoralist populations are grouped together with peripatetic communities that share their cultural traits (Figs. S6 and 3b), suggesting that migration of NRY lineages follows a path that is similar to horizontally transmitted cultural features.

Taken together, our results highlight the importance of the matrilineal rule of descent in shaping sex-specific patterns of population diversity and differentiation, stressing the need to better understand how regularities disclosed at the global level are associated with demographic processes occurring at local scales.

## Acknowledgements

We thank all sample donors for their participation in this study, the governments of Namibe and Kunene Provinces in Angola for supporting our work, João Guerra, Raimundo Dungulo, and Serafim Nemésio for assistance in the preparation of field work, António Mbeape, José Domingos, and Okongo Toko for assistance with sample collection, Roland Schröder for assistance in the lab and Enrico Macholdt for assistance in the processing of sequencing data. Financial support for this research was provided by FEDER funds through the Operational Programme for Competitiveness Factors— COMPETE, by National Funds through FCT—Foundation for Science and Technology under the PTDC/BIA-EVF/ 2907/2012 and FCOMP-01-0124-FEDER-028341, and by the Max Planck Society. SO was supported by the FCT grant SFRH/BD/85776/2012. BP acknowledges the LABEX ASLAN (ANR-10-LABX-0081) of Université de Lyon for its financial support within the program “Investissements d’Avenir” (ANR-11-IDEX-0007) of the French government operated by the National Research Agency (ANR). This is scientific paper no. 8 from the Portuguese-Angolan TwinLab established between CIBIO/InBIO and ISCED/Huíla, Lubango. This work is dedicated to the memory of our colleague Samuel Aço.

## Author contributions

JR, BP, and MS planed the study; JR, AMF, SO, TA, and FL performed the fieldwork. SO generated the laboratory data. SO, AH, BP, MS, and JR analyzed the data. SO and JR wrote the article with input from all other authors.

## References

1 Webster TH, Sayres MAW. Genomic signatures of sex-biased demography: progress and prospects. Curr Opin Genet Dev 2016; 41: 62–71.

2 Heyer E, Chaix R, Pavard S, Austerlitz F. Sex-specific demographic behaviours that shape human genomic variation. Mol Ecol 2012; 21: 597–612.

3 Jobling MA, Tyler-Smith C. Human Y-chromosome variation in the genome-sequencing era. Nat Rev Genet 2017; 18: 485–497.

4 Karmin M, Saag L, Vicente M et al. A recent bottleneck of Y chromosome diversity coincides with a global change in culture. Genome Res 2015; 25: 459–466.

5 Hallast P, Batini C, Zadik D et al. The Y-chromosome tree bursts into leaf: 13,000 high-confidence SNPs covering the majority of known clades. Mol Biol Evol 2014; 32: 661–673.

6 Lippold S, Xu H, Ko A et al. Human paternal and maternal demographic histories: Insights from high-resolution Y chromosome and mtDNA sequences. Investig Genet 2014; 5: 13.

7 Poznik GD, Xue Y, Mendez FL et al. Punctuated bursts in human male demography inferred from 1,244 worldwide Y-chromosome sequences. Nat Genet 2016; 48: 593–599.

8 Kumar V, Langstieh BT, Madhavi K V. et al. Global patterns in human mitochondrial DNA and Y-chromosome variation caused by spatial instability of the local cultural processes. PLoS Genet 2006; 2: 420–424.

9 Oliveira S, Fehn AM, Aço T et al. Matriclans shape populations: Insights from the Angolan Namib Desert into the maternal genetic history of southern Africa. Am J Phys Anthropol 2018; 165: 518–535.

10 Barnard A. Hunters and Herders of Southern Africa: A Comparative Ethnography of the Khoisan Peoples. Cambridge University Press: Cambridge, 1992.

11 Estermann C. The Ethnography of Southwestern Angola: The Herero People (translated and edited by Gibson GD). Africana Pub. Co, 1981.

12 Berland JC, Rao A. Customary Strangers:new perspectives on peripatetic people in the Middle East, Africa, and Asia. Greenwood Publishing Group, 2004.

13 MacCalman H, Grobbelaar B. Preliminary report of two stone-working OvaTjimba groups in the northern Kaokoveld of South West Africa. Staatsmuseum: Windhoek, 1965.

14 Estermann C. The Ethnography of Southwestern Angola: The Non-Bantu Peoples. The Ambo Ethnic Group (translated and edited by Gibson GD). Africana Pub. Co, 1976.

15 Güldemann T. A linguist’s view: Khoe-Kwadi speakers as the earliest food-producers of southern Africa. South African Humanit 2008; 20: 93–132.

16 Pinto JC, Oliveira S, Teixeira S et al. Food and pathogen adaptations in the Angolan Namib desert: Tracing the spread of lactase persistence and human African trypanosomiasis resistance into southwestern Africa. Am J Phys Anthropol 2016; 161: 436–447.

17 Kutanan W, Kampuansai J, Changmai P et al. Contrasting maternal and paternal genetic variation of hunter-gatherer groups in Thailand. Sci Rep 2018; 8: 1536.

18 Renaud G, Stenzel U, Kelso J. LeeHom: Adaptor trimming and merging for Illumina sequencing reads. Nucleic Acids Res 2014; 42: e141.

19 Renaud G, Stenzel U, Maricic T, Wiebe V, Kelso J. DeML: Robust demultiplexing of Illumina sequences using a likelihood-based approach. Bioinformatics 2015; 31: 770–772.

20 Poznik GD. Identifying Y-chromosome haplogroups in arbitrarily large samples of sequenced or genotyped men. bioRxiv 2016; 88716.

21 Trombetta B, D’Atanasio E, Massaia A et al. Phylogeographic Refinement and Large Scale Genotyping of Human Y Chromosome Haplogroup E Provide New Insights into the Dispersal of Early Pastoralists in the African Continent. Genome Biol Evol 2015; 7: 1940–1950.

22 Poznik GD, Henn BM, Yee MC et al. Sequencing Y chromosomes resolves discrepancy in time to common ancestor of males versus females. Science 2013; 341: 562–565.

23 Excoffier L, Lischer HEL. Arlequin suite ver 3.5: A new series of programs to perform population genetics analyses under Linux and Windows. Mol Ecol Resour 2010; 10: 564–567.

24 Drummond AJ, Suchard MA, Xie D, Rambaut A. Bayesian phylogenetics with BEAUti and the BEAST 1.7. Mol Biol Evol 2012; 29: 1969–1973.

25 Barbieri C, Hübner A, Macholdt E et al. Refining the Y chromosome phylogeny with southern African sequences. Hum Genet 2016; 135: 541–553.

26 Mallick S, Li H, Lipson M et al. The Simons Genome Diversity Project: 300 genomes from 142 diverse populations. Nature 2016; 538: 201–206.

27 Browning SR, Browning BL. Rapid and Accurate Haplotype Phasing and Missing-Data Inference for Whole-Genome Association Studies By Use of Localized Haplotype Clustering. Am J Hum Genet 2007; 81: 1084–1097.

28 Scozzari R, Massaia A, Trombetta B et al. An unbiased resource of novel SNP markers provides a new chronology for the human y chromosome and reveals a deep phylogenetic structure in Africa. Genome Res 2014; 24: 535–544.

29 Henn BM, Gignoux C, Lin AA et al. Y-chromosomal evidence of a pastoralist migration through Tanzania to southern Africa. Proc Natl Acad Sci 2008; 105: 10693–10698.

30 Beleza S, Gusmão L, Amorim A, Carracedo A, Salas A. The genetic legacy of western Bantu migrations. Hum Genet 2005; 117: 366–75.

31 De Filippo C, Barbieri C, Whitten M et al. Y-chromosomal variation in sub-Saharan Africa: Insights into the history of Niger-Congo groups. Mol Biol Evol 2011; 28: 1255–1269.

32 Fenner JN. Cross-cultural estimation of the human generation interval for use in genetics-based population divergence studies. Am J Phys Anthropol 2005; 128: 415–423.

33 Bajić V, Barbieri C, Hübner A et al. Genetic structure and sex-biased gene flow in the history of southern African populations. bioRxiv 2017; 237297.

34 Heller R, Chikhi L, Siegismund HR. The Confounding Effect of Population Structure on Bayesian Skyline Plot Inferences of Demographic History. PLoS One 2013; 8: e62992.

35 Schlebusch CM, Malmström H, Günther T et al. Southern African ancient genomes estimate modern human divergence to 350,000 to 260,000 years ago. Science 2017; 358: 652–655.

36 Rocha J, Fehn A-M. Genetics and Demographic History of the Bantu. eLS 2016.

37 Verdu P, Becker NSA, Froment A et al. Sociocultural behavior, sex-biased admixture, and effective population sizes in central African pygmies and non-pygmies. Mol Biol Evol 2013; 30: 918–937.

38 Destro-Bisol G, Donati F, Coia V et al. Variation of female and male lineages in sub-Saharan populations: The importance of sociocultural factors. Mol Biol Evol 2004; 21: 1673–1682.

39 Barbieri C, Vicente M, Oliveira S et al. Migration and interaction in a contact zone: mtDNA variation among Bantu-speakers in Southern Africa. PLoS One 2014; 9: e99117.

40 Wood ET, Stover DA, Ehret C et al. Contrasting patterns of Y chromosome and mtDNA variation in Africa: Evidence for sex-biased demographic processes. Eur J Hum Genet 2005; 13: 867–876.

41 Scelza BA.Female choice and extra-pair paternity in a traditional human population. Biol Lett 2011; 7: 889–891.

42 Chaix R, Quintana-Murci L, Hegay T et al. From social to genetic structures in central Asia. Curr Biol 2007; 17: 43–48.

43 Hartl D, Clark A. Principles of population genetics. Sunderland: Sinauer associates, 1997.

44 Zeng TC, Aw AJ, Feldman MW. Cultural hitchhiking and competition between patrilineal kin groups explain the post-Neolithic Y-chromosome bottleneck. Nat Commun 2018; 9: 2077.

45 Jakobsson M, Edge MD, Rosenberg NA. The relationship between F(ST) and the frequency of the most frequent allele. Genetics 2013; 193: 515–28.

46 Fix A. Migration and colonization in human microevolution. Cambridge University Press: Cambridge, 1999.

